# SplicedFamAlign: CDS-to-gene spliced alignment and identification of transcript orthology groups

**DOI:** 10.1101/420307

**Authors:** Safa Jammali, Jean-David Aguilar, Esaie Kuitche, Aïda Ouangraoua

## Abstract

**Motivation:** The inference of splicing orthology relationships between gene transcripts is a basic step for the prediction of transcripts and the annotation of gene structures in genomes. Spliced alignment that consists in aligning a spliced cDNA sequence against an unspliced genomic sequence, constitutes a promising, yet unexplored approach for the identification of splicing orthology relationships. Existing spliced alignment algorithms do not exploit the information on the splicing structure of the input sequences, namely the exon structure of the cDNA sequence and the exon-intron structure of the genomic sequences. Yet, this information is often available for coding DNA sequences (CDS) and gene sequences annotated in databases, and it can help improve the accuracy of the computed spliced alignments. To address this issue, we introduce a new spliced alignment problem and a method called SplicedFamAlign (SFA) for computing the alignment of a spliced CDS against a gene sequence while accounting for the splicing structures of the input sequences, and then the inference of transcript splicing orthology groups in a gene family based on spliced alignments.

**Results:** The experimental results show that SFA outperforms existing spliced alignment methods in terms of accuracy and execution time for CDS-to-gene alignment. We also show that the performance of SFA remains high for various levels of sequence similarity between input sequences, thanks to accounting for the splicing structure of the input sequences. It is important to notice that unlike all current spliced alignment methods that are meant for cDNA-to-genome alignments and can be used for CDS-to-gene alignments, SFA is the first method specifically designed for CDS-to-gene alignments. We show its usefulness for the comparison of genes and transcripts within a gene family for the purpose of analyzing splicing orthologies. It can also be used for gene structure annotation and alternative splicing analyses.

**Availability:** SplicedFamAlign was implemented in Python. Source code is freely available at https://github.com/UdeS-CoBIUS/SpliceFamAlign

## 1 Introduction

A spliced alignment is an alignment of a partial or full length spliced transcript sequence against an unspliced genomic sequence [8]. A spliced alignment allows to highlight the boundaries and the alignment of exons of the transcript sequence on the genomic sequence. It is an effective method for gene recognition, gene structure prediction and alternative splicing analyses [2, 11, 15, 19, 21].

Several methods have been developed to address different versions of the cDNA-to-genome spliced alignment problem, which consists in finding an optimal alignment of a spliced cDNA sequence against an unspliced genomic sequence, given an optimal function [3, 9, 12, 14, 23, 25]. A complete overview of these methods is provided in [25] and complemented by [12]. These methods follow a general scheme that consists in first identifying candidate target alignment locations on the genomic sequence for thecDNAsequence, and then computing approximate spliced alignments of the cDNA sequence against each candidate target genomic regions. For computing approximate spliced alignments, the first criterion of optimality used by all spliced alignment methods is the sequence similarity. In addition to sequence similarity, some methods also account for splice signals on the unspliced genomic sequence such as canonical dinucleotide splice signals “GT” and “AG” at extremities of an intron, in their criterion of optimality in order to infer accurate exon boundaries in the alignments. It has been shown that the performance of methods accounting for splice signals is superior to that of only sequence similarity-based methods [12, 25]. Indeed, splice signals constitute strong structural signals that are used by splice signal-based methods for inferring accurate splice sites. Thus, the use of splice signals explains the superiority of splice signal-based methods over only sequence similarity-based methods. However, none of the existing spliced alignment methods takes into account the splicing structure of the input sequences, namely the exon structure of the cDNA sequence and the exon-intron structure of the genomic sequences, in addition to splice signals and sequence similarity. Yet, the information on the splicing structure and known splice sites is often available in annotation of CDS and gene sequences. This information can be used to improve the accuracy of spliced alignments.

In this paper, we re-visit the spliced alignment pro of computing accurate CDS-to-gene spliced alignm transcript orthology groups within a set of transcr gene family. Identifying orthologous isoforms at tran requisite to describing evolutionary relationships bet of splicing structure and sets of splice variants [5, 17]. Here, we focus on the spliced alignment of full CDS against gene sequences within a gene family, which allows to identify splicing orthologous CDS with similar sequences and splicing structures from genes that have evolved from a common ancestral gene [26, 1]. Splicing orthologs are supposed to have retained the same function in the course of evolution. Identifying splicing orthologs using spliced alignments requires precise alignment of exons and location of their boundaries in the spliced alignments. To achieve this aim, we introduce a new version of the spliced alignment problem that accounts for the splicing structure of the input sequences. For this version of the problem, we propose SplicedFamAlign (SFA), a method for fast and accurate alignment of spliced CDS against unspliced gene sequences (*SFA-align*), and for the identification of splicing orthologs using spliced alignments (*SFA-ortholog*).

The SFA-align algorithm (see overview in Figure 1) starts by fastly computing local alignments in order to identify highly conserved sequences between the input CDS and gene sequences. These local alignments are used as anchors and each anchor alignment is trimmed at the extremities in order to cover at most one exon on the CDS. Next, a gapped extension algorithm accounting for CDS exon boundaries is used to extend anchor alignments in both direction in order to maximize the coverage of CDS exons. Finally a global spliced alignment algorithm accounting for the exon boundaries of the CDS and the gene sequence can be applied for aligning the remaining segments between the extended anchors. This anchored spliced alignment approach has been used by several other methods such as Splign [12], Spidey [24] and MGAlign [18]. The added value of the SFA method is that it makes use of the splicing structure of the input sequences in its local and global spliced alignment steps, in addition to the splice signals on the genomic sequence, in order to produce more accurate spliced alignments.

**Fig. 1:**
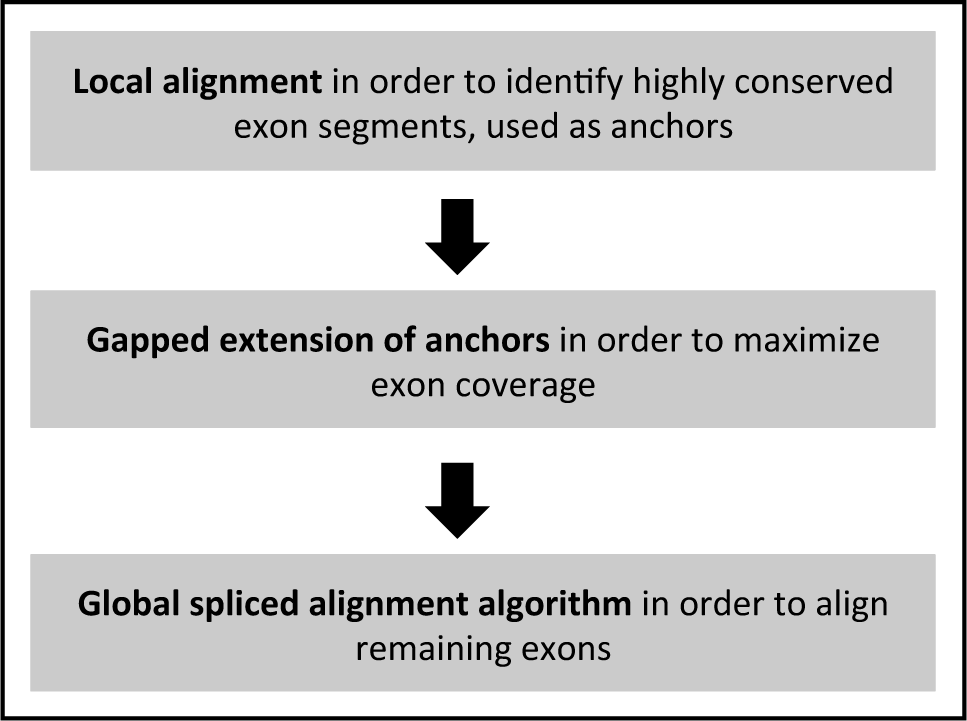
Overview of the SFA-align algorithm.

The SFA-ortholog algorithm starts by identifying pairwise orthologous CDS. The definition of the orthology relation between two CDS relies on the preservation of their splicing structures. We say that a spliced alignment of a CDS sequence against a gene sequence preserves the exon structure of the CDS, if the spliced alignment induces a sequence conservation of all the exons of the CDS on the gene sequence, and it also induces an intron between any two consecutive exons of the CDS, and no intron within an exon of the CDS (see Figure 2 for example in which the exon structures of both CDS1 and CDS2 are preserved by their spliced alignments against a gene sequence).

**Fig. 2:**
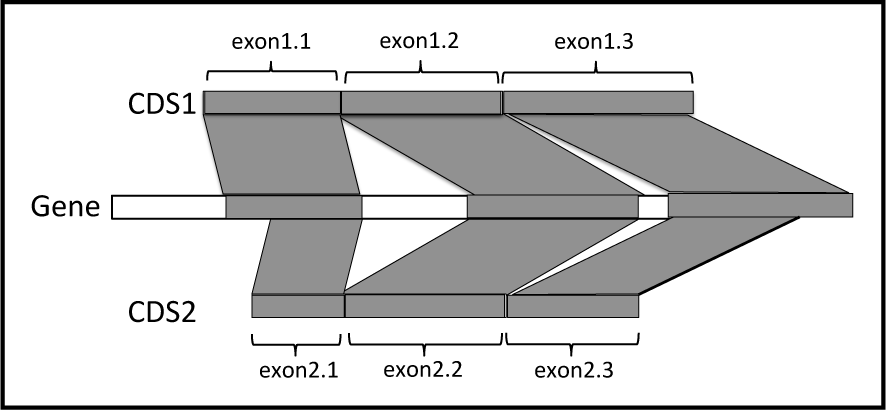
Two orthologous CDS, according to the definition of splicing orthology: a) the exon structures of the two CDS are preserved by their splicing alignments against the gene sequence and b) the induced one-to-one correspondence between the exons of CDS1 and CDS2 is (exon1.1,exon2.1),(exon1.2,exon2.2), (exon1.3,exon2.3).

Next, two CDS are *splicing orthologs* if their spliced alignments against the gene of one of the two CDS a) preserve their exon structures, and b) induce a one-to-one correspondence between the exons of the two CDS (see Figure 2 for example). Finally, the pairs of orthologous CDS within a gene family are used to define an orthology graph whose nodes are all the CDS of the family and edges represent pairwise orthology relations between the CDS. Then, assuming perfect pairwise orthology relations,the CDS orthology groups are defined as the connected components of the orthology graph.

The main contributions of SplicedFamAlign can be described as follows:

1. It allows fast computing of accurate CDS-to-gene spliced alignments, and accurate CDS splicing orthology groups within a gene family.
2. Its performance remains high for various levels of sequence similarity, thanks to the use of the splicing structure of input sequences, which allows to detect splicing structure conservation even in the cases of low sequence conservation.
3. In the case where the splicing structure of sequences are not given as input, SplicedFamAlign includes a preliminary step that allows computing the splicing structure of input sequences by aligning each CDS against its own gene.

## 2. Results and discussion

### 2.1 Dataset

SplicedFamAlign was evaluated based on a dataset of three real sets of homologous genes and three simulated sets of gene families.

#### 2.1.1 Real data

The dataset contains homologous genes with their CDS sequences from 3 gene families FAM86, MAG and TP53 from the Ensembl-Compara database release 85 [6]. For each family, 8 homologous genes were selected with 14 CDS for FAM86, 26 for MAG and 51 for TP53. For each set of homologous genes, the genes are from 6 different amniote species, *human, chimpanzee, mouse, rat, cow and chicken* (for FAM86 and TP53) or *lizard* (for MAG). Table 1 gives more details about the real dataset.

**Table 1.** Detailed description of the real data, from the Ensembl-Comapara database, used for the evaluation. For each gene family, the following information are given: the species name, the Ensembl identifier of gene, the number of CDS for each gene, the average CDS length, the average gene length, and the average pairwise Percent Sequence Identity (PID). The average pairwise PID were computed based on pairwise alignments of the CDS obtained from the multiple alignments of their proteins families provided by Ensembl-Comapara.

| Species | Family |  |  |  |  |  |
| --- | --- | --- | --- | --- | --- | --- |
|  | FAM86 |  | MAG |  | TP53 |  |
|  | Gene_ID | #CDS | Gene_ID | #CDS | Gene_ID | #CDS |
| Human | ENSG00000158483 | 3 | ENSG00000105492 | 6 | ENSG00000141510 | 15 |
|  | ENSG00000186523 | 4 | ENSG00000142512 | 7 | ENSG00000073282 | 11 |
|  | ENSG00000145002 | 2 | ENSG00000105695 | 4 | ENSG00000078900 | 9 |
| Chimpanzee | ENSPTRG00000007738 | 1 | ENSPTRG00000011374 | 1 | ENSPTRG00000008703 | 1 |
| Mouse | ENSMUSG00000022544 | 1 | ENSMUSG00000051504 | 4 | ENSMUSG00000022510 | 8 |
| Rat | ENSRNOG00000002876 | 1 | ENSRNOG00000021023 | 2 | ENSRNOG00000010756 | 4 |
| Cow | ENSBTAG00000008222 | 1 | ENSBTAG00000017044 | 1 | ENSBTAG00000001069 | 1 |
| Chicken | ENSGALG00000002044 | 1 |  |  | ENSGALG00000007324 | 2 |
| Lizard |  |  | ENSACAG00000005408 | 1 |  |  |
| Total | 8 genes | 14 | 8 genes | 26 | 8 genes | 51 |
| Avg. CDS length | 726 |  | 1397.42 |  | 1277.41 |  |
| Avg. gene lengths | 22782.37 |  | 26049.75 |  | 109457.75 |  |
| Avg. pairwise PID | 56.57 |  | 41.41 |  | 57.21 |  |

FAM86 is a group of genes with sequence similarity and unknown function. MAG is the family of Myelin Associated Glycoprotein genes whose proteins are type I membrane protein and member of the immunoglobulin superfamily. TP53 is a family of genes encoding for tumor suppressor proteins p53.

#### 2.1.2 Simulated Data

We generated a dataset of simulated gene families with gene and cDNA sequences accounting for gene splicing structure evolution and alternative splicing events. We designed a new simulation method that takes as input a gene tree with branch lengths representing evolutionary rates, generates an ancestral gene at the root of the tree, and makes this gene evolve along branches of the gene tree. The ancestral root gene is generated with an exon-intron structure and an initial set of CDS of alternative transcripts for the gene, based on parameters learned from a dataset of 10.000 vertebrate genes from amniote species from the Ensembl database [6]. The evolution simulated along branches of the tree accounts for two levels of evolution. First, at the level of genes, the following evolutionary events acting on the splicing structure of genes are included, exon duplication, exon gain and exon loss. Evolutionary events acting on the sequence of coding exons, namely nucleotide insertion, deletion and substitution events are also included at the level of genes, using empirical codon evolution models [16]. Second, at the level of transcripts, we account for two events acting on the sets of transcripts and CDS generated by genes, isoform creation, and isoform loss. The isoform creation event corresponds to the acquisition by a gene of the ability to produce a new transcript through a new combination of exons. The isoform loss event corresponds to the loss of the ability to produce a given transcript.

Using the simulation method, we generated three sets of gene family, a first set called *Small* for which the evolutionary rates on the gene tree branches are low, a second set called *Medium* with medium evolutionary rate, and a third set called *Large* with a high evolutionary rates. Each of the three set contains 36 simulated gene families with 5 genes and 5 to 17 CDS in total. For each family in each set, we generated a set of 5 gene sequences with their CDS, all true pairwise spliced alignments between any CDS and any gene of the family, the true multiple sequence alignment of all CDS and gene sequences, and all true splicing orthology relations between CDS. Table 2 gives more details about the simulated datasets used for the evaluation.

**Table 2.** Detailed description of the simulated data used for the evaluation: For each simulated dataset, the number of families, the number of genes per family, the average number of CDS, the average CDS length, the average gene length, and the average pairwise PID in families are given. For the average measures, the standard deviations are also given.

|  | Small | Medium | Large |
| --- | --- | --- | --- |
| Number of families | 36 | 36 | 36 |
| Number of genes per family | 5 | 5 | 5 |
| Avg. number of CDS | 8.86<br>(2.95) | 8.03<br>(2.78) | 9.30<br>(2.87) |
| Avg. CDS length | 808.24<br>(175.79) | 779.44<br>(186.66) | 799.40<br>(214.40) |
| Avg. gene length | 2556.83<br>(537.93) | 2358.60<br>(548.06) | 2477.95<br>(552.02) |
| Avg. pairwise PID | 76.25<br>(18.92) | 39.20<br>(13.85) | 11.42<br>(3.86) |

### 2.2 Evaluated methods

SplicedFamAlign results were compared with the results of Splign [12], the most recent and current best performing cDNA-to-genome spliced alignment method. In [12], Splign was compared with SIM4 [7], Spidey [24], BLAT [14], GMAP [25], and Spa [23] in terms of the capacity of the method to realize alignments at various levels of similarity between input sequences. Splign was shown to be more performant that other methods in the comparison for all levels of sequence similarity. Three version of SplicedFamAlign were tested. SFA_L corresponds to the first local alignment step of SplicedFamAlign. SFA_E is SFA_L followed by the gapped extension step of the method. SFA_G is SFA_E followed by the final global alignment step of the method. A description of the methods used in the evaluation is given in Table 3.

**Table 3.**
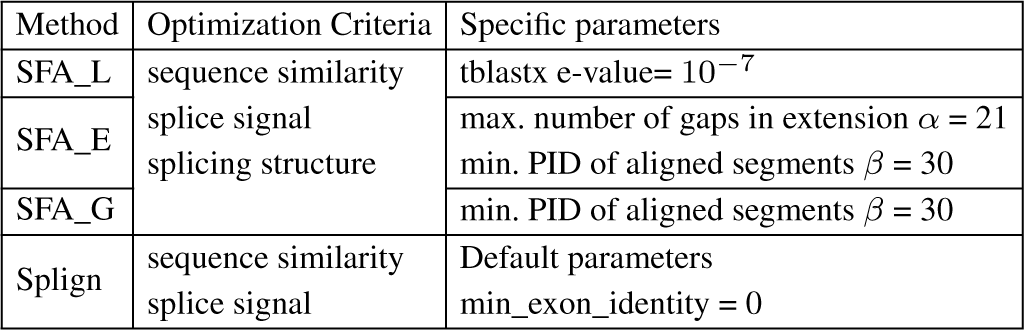
Description of the methods used in the evaluation: For each method, the optimization criteria and the specific parameters are given.

For SFA_E, the maximum number of gaps in a left or right extension of an anchor alignment is *α* = 21. For SFA_E and SFA_G, the additional aligned segments obtained after the anchor extension step or the global alignment step are kept if and only if they have a Percent Sequence Identity (PID) greater or equal to *β* = 30% (See Section 3.3 for a more detailed description of the parameters of algorithms used in the anchor extension and the global alignment steps of SFA).

For Splign, the parameter min_exon_identity is set to 0 in order to allow Splign achieve the highest coverage of CDS possible. Setting this parameter to higher values makes Splign get worse performance (data not shown).

### 2.3 Discussion

First, we compared the ability of the methods to compute spliced alignments with high coverage of CDS (Figure 3), relevant Percent Sequence Identity (PID) (Figure 4), and induced exon extremities corresponding to actual exon extremities in the CDS and the gene sequence (Figure 5). For the simulated data, for which the true alignment of the sequences in provided, we compared the ability of the methods to recover the true spliced alignments (Figure 6).

**Fig. 3:**
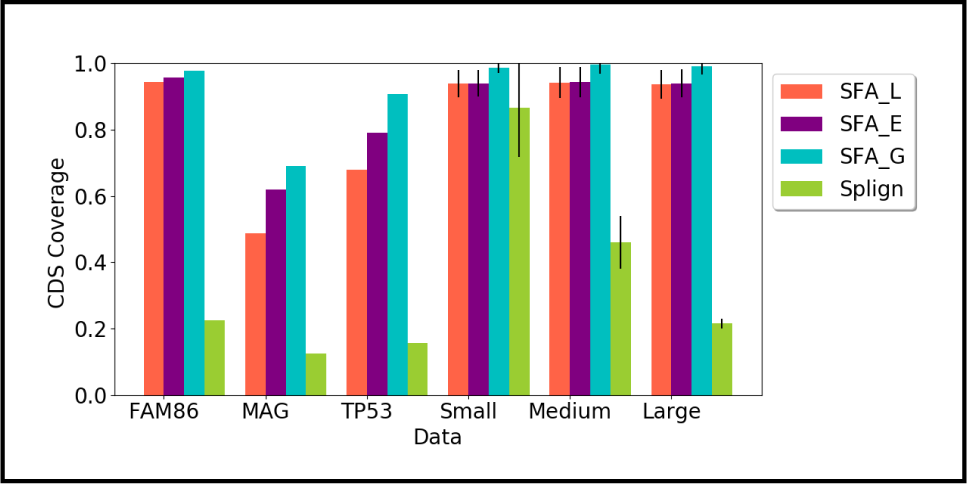
Average CDS coverage of spliced alignments obtaining using the methods (SFA_L, SFA_E, SFA_G, Splign) on the real and simulated dataset. For the simulated datasets composed of several gene families, the standard deviations are also given.

**Fig. 4:**
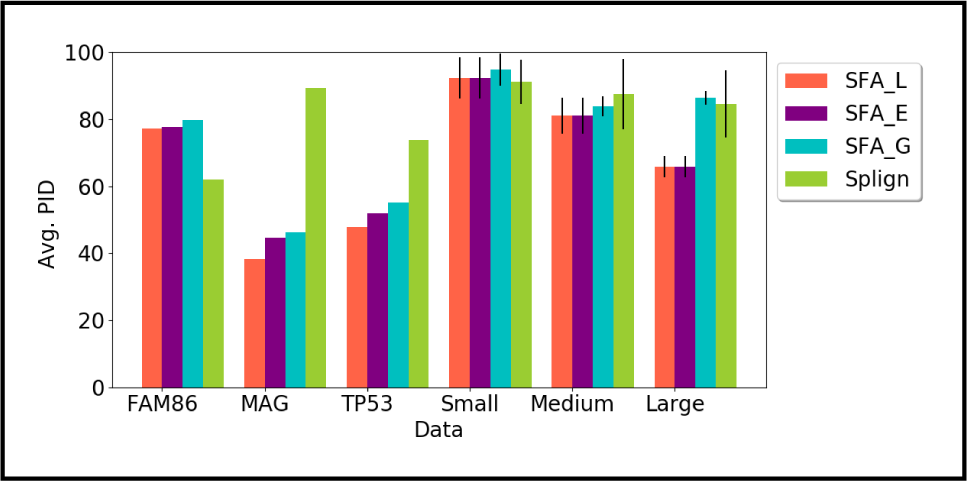
Average Percent Sequence Identity (PID) for each method and each dataset used in the evaluation. For the simulated datasets composed of several gene families, the standard deviation are also given.

**Fig. 5:**
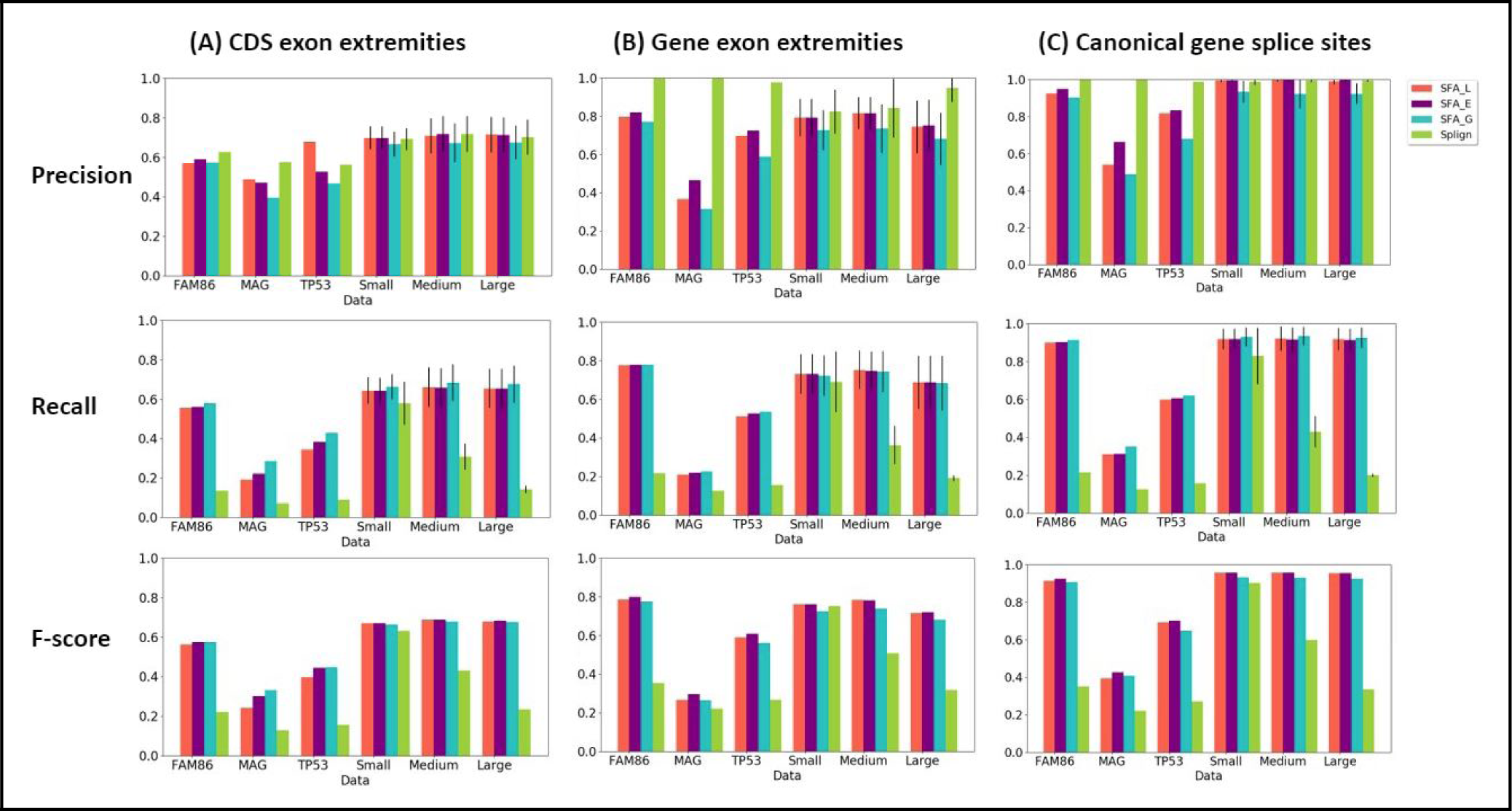
Precision, recall and f-score measure, for all methods (SFA_L, SFA_E, and SFA_G, Splign), for the comparison of inferred exon extremities with real CDS exon extremities (A), real gene exon extremities (B), canonical gene splice sites (C). For the simulated datasets composed of several gene families, the standard deviation are also given for precision and recall.

**Fig. 6:**
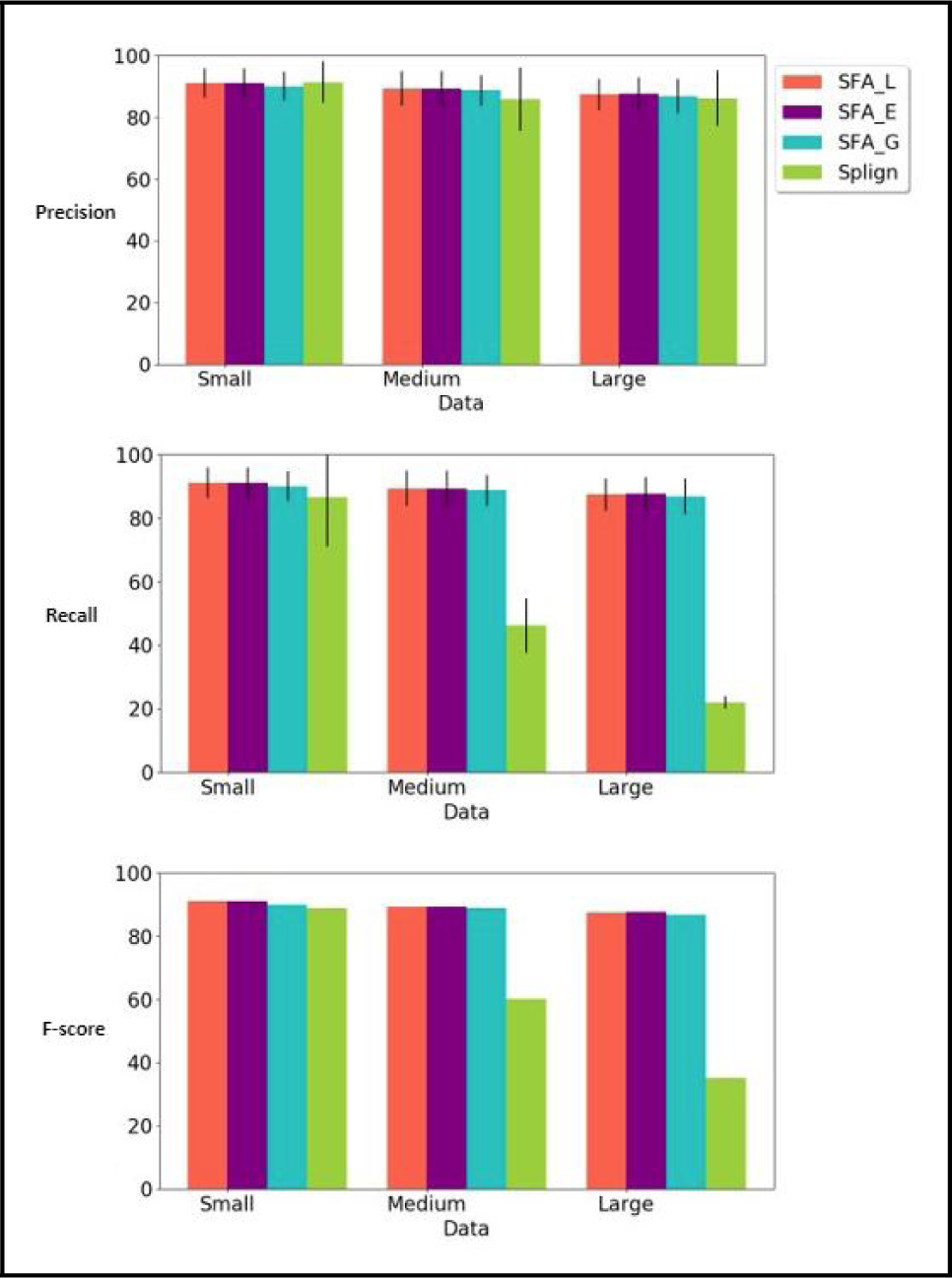
Precision, recall and f-score measure, for all methods (SFA_L, SFA_E, and SFA_G, Splign), for the comparison of the computed spliced alignments with the true spliced alignments. For each simulated dataset, the standard deviation are also given for precision and recall.

Second, the true orthology relationships between the CDS are also known for the simulated data. We compared the ability of the methods to be used for computing the true CDS orthology groups within a gene family based on spliced alignments (Figure 7).

**Fig. 7:**
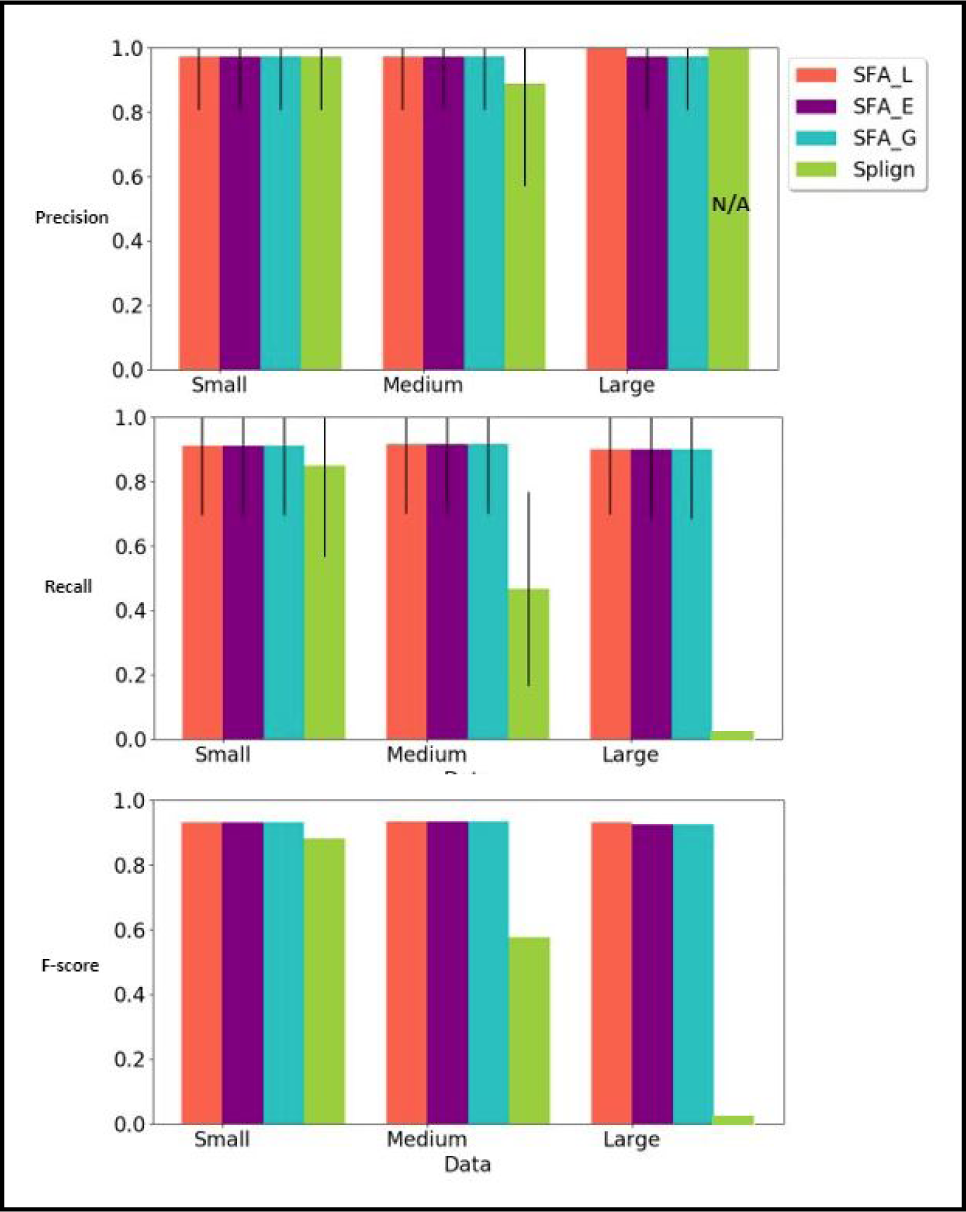
Precision, recall and f-score measures, for all methods (SFA_L, SFA_E, and SFA_G, Splign), for the comparison of the computed CDS orthology groups with the true CDS orthology groups. For each simulated dataset, the standard deviation are also given for precision and recall.

Third, we compared the execution time of the methods (Figure 8).

**Fig. 8:**
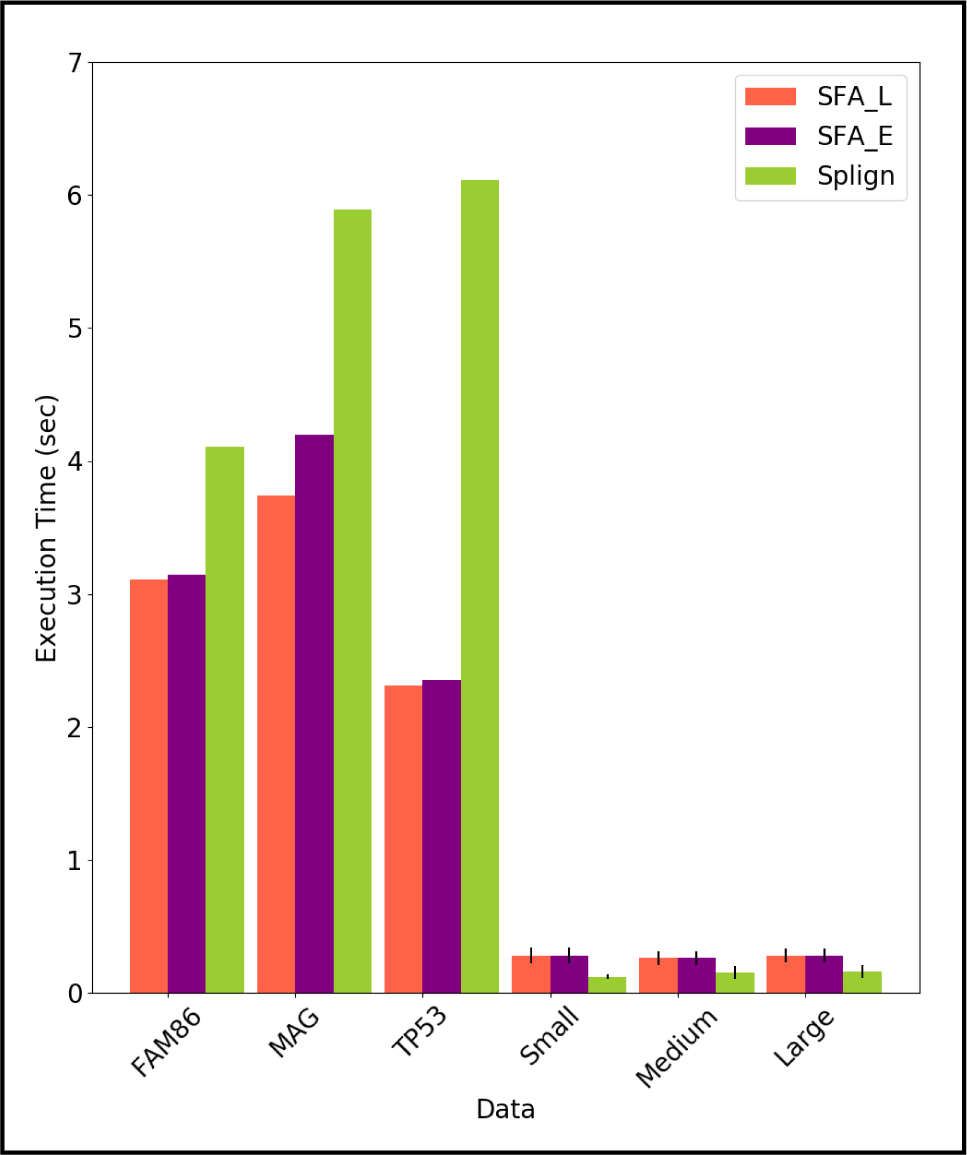
Average execution times in second (sec) of SFA_L, SFA_E and Splign to compute spliced alignments for all datasets. For each simulated dataset, the standard deviation are also given.

#### 2.3.1 Evaluation of the quality of spliced alignments

**CDS coverage**. We compared the CDS coverage of the spliced alignments computed using the methods. The results are shown in Figure 3. In terms of CDS coverage, SFA methods and especially SFA_G, show the best results and their performances remain high for various levels of similarity between input sequences, covering in average more than 90% of CDS for all simulated datasets (Small, Medium, Large). Splign achieves a high CDS coverage when the sequences have a high level of similarity (Small dataset), but the performance decreases for medium and low similarity levels.

**Percent Sequence Identity (PID)**. The PID of the spliced alignments constitutes also a good criterion to evaluate the quality of the alignments. For all methods and all datasets, the PID computed for their spliced alignment results are shown in Figure 4. The PID for SFA_L and SFA_E decreases when the input sequence similarity decreases, which is expected. However, for the other methods, especially Splign, the PID remains high for most datasets (more than 80%). For instance, while Splign achieves the lowest CDS coverage for the MAG family (Figure 3), it achieves the highest PID for this family. Taken together, the results on the CDS coverage and the PID (Figure 3 and 4) suggest that Splign is very stringent and aligns only highly sequence-conserved segments, whereas the SFA methods align also less similar segments.

**Inference of real exon extremities**. We compared the ability of the methods to correctly identify actual exon extremities in the CDS and the gene sequence and canonical splice sites in the gene sequence. We used the following performance metrics. The precision measure represents the fraction of inferred exon extremities that corresponds to real CDS exon extremities (A), real gene exon extremities (B), canonical gene splice sites (C). The recall measure represents the fraction of real CDS exon extremities (A), real gene exon extremities (B), canonical gene splice sites (C) that are correctly inferred by the spliced alignments. The f-score is the harmonic mean of precision and recall.

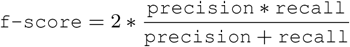

The results of the comparison are shown in Figure 5. In terms of precision, we observe that, for all references (A), (B) and (C), all methods achieve a good performance. Globally, they perform better for gene exon extremities (B) and canonical splice sites (C) than for CDS exon extremities (A). In particular, Splign performs almost perfectly for (B) and (C). SFA_E is the second one in terms of precision and also performs well.

In terms of recall, SFA_G is the best performing method for all references (A), (B) and (C), closely followed by SFA_E and SFA_L. The performance of SFA methods remains similar for various levels of sequence similarity (Small, Medium and Large datasets). However, the performance of Splign decreases when the level of sequence similarity decreases. This observation is coherent with the results obtained for the evaluation of the CDS coverage achieved by the methods. Indeed, the lower the CDS coverage, the lower the recall measure achieved.

Combining the precision and the recall measures, the f-score measure shows that SFA_E and SFA_L are the best performing methods. The robustness of the SFA methods to changes in the level of sequence similarity can be explained by the use of the splicing structure of the input sequences that allows to detect structure conservation even when the sequence conservation signal is low.

**Comparison with true alignments**. For the simulated datasets for which the true spliced alignments of CDS against gene sequences are provided, we evaluated the ability of the methods to correctly recover the true spliced alignments. We evaluated the precision, recall and f-score measure of a computed spliced alignment as follows. The precision measure is the fraction of pairs of aligned nucleotides in the computed alignment that are also aligned together in the true alignment. The recall measure is the fraction of pairs of aligned nucleotides in the true alignment that are also aligned together in the computed alignment.

The f-score is a combination of the precision and recall as defined for the evaluation of the ability to infer real exon extremities.

The results of the evaluation are shown in Figure 6. We can observe that all methods achieve high and comparable precision rates. In terms of recall, the SFA methods also have a high performance for various levels of sequence similarity. Splign also performs very well for high sequence similarities (Small dataset), but the performance decreases with the decrease of sequence similarity.

#### 2.3.2 Evaluation of the quality for CDS orthology groups identification

We applied our algorithm for the identification of CDS orthology groups based on structural similarity, using spliced alignments computed by the methods SFA_L, SFA_E, and SFA_G and Splign. For each method, we then obtained a set of CDS orthology groups.

We evaluated and compared the ability of the methods to recover the true CDS orthology groups. Two CDS are true orthologs if the sets of exons composing them are in bijection in such a way that any pair of exons in bijection descend from a same ancestral exon. Thus, we compared the CDS orthology groups obtained using each method with the true CDS orthology groups given in the simulated datasets. The precision, recall and f-score measures of a computed clustering are defined as follows.

The precision represents the fraction of pairs of CDS found as orthologs by the computed clustering that are true orthologs. The recall represents the fraction of true pairs of orthologous CDS that are also found as orthologs by the computed clustering. The f-score is a combination of the precision and recall as defined in Section 2.3.1.

Figure 7 shows the results for each method. The precision score for Splign on the dataset Large is N/A because no pair of CDS were found as orthologs by the computed clustering. As for previous comparisons, the precision scores are high for all methods. For the SFA methods, the recall scores are also high and robust to changes in the level of sequence similarity, whereas for Splign, the recall score decreases when the level of sequence similarity decreases.

#### 2.3.3 Evaluation of the execution time

Finally, we compared the execution time of the methods. The average execution times of the SFA_L, SFA_E and Splign methods to compute spliced alignments for each dataset are shown in Figure 8. The execution times of SFA_G are not displayed in the same figure as they are more than 500 times higher than the execution times of other methods. The very high execution times of SFA_G are explained by the global alignment step of the method that uses a dynamic programming algorithm of quadratic time complexity with a multiplicative constant *I*_*max*_ *- I*_*min*_ = 5000 (See Section 3.3, Step 3 for a description of the main recurrence formula used by the dynamic programming algorithm). For instance, in contrast to Splign which stops without achieving its global alignment step when the sequence similarity is low, the global alignment step of SFA_G is achieved even if the preceding steps of local alignment and anchor extension end up with a spliced alignment having a low CDS coverage. For the remaining methods, the execution times of SFA_E are slightly higher than those of SFA_L. Splign has higher execution times than SFA_L and SFA_E for real datasets, but slightly lower execution times for simulated datasets. This can be explained by the smaller difference between CDS lengths and gene lengths in the simulated datasets compared to the real data. (See the average CDS and gene lengths of datasets in Tables 1 and 2). So, when the time-consuming global alignment step of Splign is achieved, it is applied on larger instances in the real dataset than in the simulated dataset.

## 3 Methods

In this section, we first give some formal definitions that will be useful for the remaining of the section. In the second subsection, the definitions of three versions of the spliced alignment problem are given under a unified framework allowing to compare theoretically the optimization criteria of the different versions of the problem. All existing cDNA-to-genome spliced alignment methods correspond either to the first or the second version of the problem that account for sequence similarity and splice signals in the input genomic sequence. The last version of the problems introduced in this paper additionally takes into account the splicing structure of the input sequences. In the third subsection, we describe our SFA-align algorithm for the new version of the spliced alignment problem. In the fourth subsection, we describe SFA-ortholog algorithm for the computation of CDS orthology groups, using spliced alignments.

### 3.1 Preliminary definitions: gene, CDS, spliced alignment

Given a set *S, |S* | denotes the size of *S*, and given a sequence *T*, length(*T*) denotes the length of *T*. Note that in the nomenclature used in this section, we call *CDS* the exon structure of a coding sequence and *CDS sequence* the nucleotide sequence of the coding sequence.

**Gene and CDS:** A gene sequence is a DNA sequence on the alphabet of nucleotides Σ = {*A, C, G, T*}. Given a gene sequence *g*, an *exon* of *g* is represented as a pair of integers (*a, b*) such that *a ≤ b* and *a* and *b* are the start and end locations of the exon on the gene sequence. The sequence of the exon (*a, b*) is then denoted by *g*[*a, b*]. A CDS of the gene sequence *g* is represented as a chain *c* = {(*a*_1_, *b*_1_), …, (*a*_*j*_, *b*_*j*_)} of exons of *g* such that for any two successive exons (*a*_*i*_, *b*_*i*_) and (*a*_*i*+1_, *b*_*i*+1_),*b*_*i*_ < *a*_*i*+1_. The i^*th*^ exon of the CDS *c* is then denoted by *c*[*i*]. The set of introns induced by the CDS *c* is denoted by Intron(*c*) =*C*{(*b*_1_, *a*_2_), (*b*_2_, *a*_3_) …, (*b*_*j*-1_, *a*_*j*_)}. The set of known CDS of a gene *g* is denoted by (*g*) and the set of all exons of all konwn CDS of *g* is denoted by *ε* (*g*) = U_*c∈C*(*g*)_ *c* (see Figure 9 for an illustration).

**Fig. 9:**
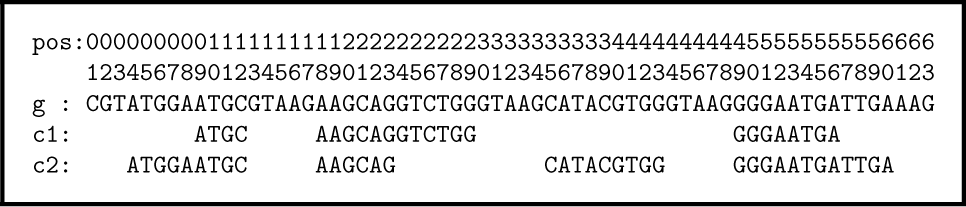
Toy example of a gene sequence *g* with nucleotides numbered by position from 1 to 63, and a set of two CDS *C*(*g*) =*{c*_1_, *c*_2_*}* such that *c*_1_ = *{*(9, 12), (18, 29), (49, 56)*}* and *c*_2_ ={(4, 12), (18, 23), (35, 43), (49, 60)}, inducing a set of exons *E* (*g*) ={(4, 12), (9, 12), (18, 23), (18, 29), (35, 43), (49, 56), (49, 60)}. The sequence of the CDS *c*_1_ is *g*[*c*_1_] =ATGCAAGCAGGTCTGGGGGAATGA with a set of exons *E* (*g*[*c*_1_]) = {(1, 4), (5, 16), (17, 24)}. The first exon of the CDS sequence *g*[*c*_1_] is (1, 4) with exon sequence *g*[*c*_1_][1, 4]=ATGC.

The sequence of a CDS *c* of *g* is denoted by *g*[*c*]. *g*[*c*] is the concatenation of the sequences of the exons composing *c*. An exon of a CDS sequence *g*[*c*] is represented as a pair of integers (*k, l*) such that *k ≤ l*, and *k* and *l* are the start and end locations of the exon on the CDS sequence. In this case, the sequence of the exon (*k, l*) of *g*[*c*] is denoted by *g*[*c*][*k, l*]. The set of exons composing a CDS sequence *g*[*c*] is denoted by *E* (*g*[*c*]) (see Figure 9 for an illustration).

**Spliced alignment:** A *spliced alignment* is an alignment of a CDS sequence against a gene sequence that allows to identify conserved exons sequences. Formally, a spliced alignment of a CDS sequence *g*[*c*] against a gene sequence *h* is represented as a chain *A* =*{*(*k*_1_, *l*_1_, *a*_1_, *b*_1_), …, (*k*_*j*_, *l*_*j*_, *a*_*j*_, *b*_*j*_)*}* of quadruplets called blocks such that for any block (*k, l, a, b*) of *A, k ≤ l* and *k* and *l* are the start and end locations of a segment on the CDS sequence, and *a ≤ b* and *a* and *b* are the start and end locations of a segment on the gene sequence or *a* = *b* = 0; The i^*th*^ block of a spliced alignment *A* is denoted by *A*[*i*], and:

1. *k*_1_ = 1, *l*_*j*_ = length(*g*[*c*]) and for any two successive blocks *A*[*i*] = (*k*_*i*_, *l*_*i*_, *a*_*i*_, *b*_*i*_) and *A*[*i* + 1] = (*k*_*i*+1_, *l*_*i*+1_, *a*_*i*+1_, *b*_*i*+1_), we have *l*_*i*_ = *k*_*i*+1_ − 1
2. for any two blocks *A*[*i*_1_] and *A*[*i*_2_] with *i*_1_ *< i*_2_, we have *b*_*i*1_ *< a*_*i*2_if *a*_*i*1_ ≠0 and *a*_*i*2_ ≠ 0.

A block (*k, l, a, b*) represents an alignment of a segment *g*[*c*][*k, l*] of the CDS sequence with a segment *h*[*a, b*] of the gene sequence. If *a* = *b* = 0, then the gene segment *h*[*a, b*] is empty and the block (*k, l, a, b*) represents a deletion of the CDS segment *g*[*c*][*k, l*] in the spliced alignment. We call such a block a *deleted block*. Otherwise, the block (*k, l, a, b*) represents a conservation between a putative CDS exon sequence *g*[*c*][*k, l*] and a putative gene exon sequence *h*[*a, b*], and we call such a block a *conserved block* (see Figure 10 for an illustration).

**Fig. 10:**
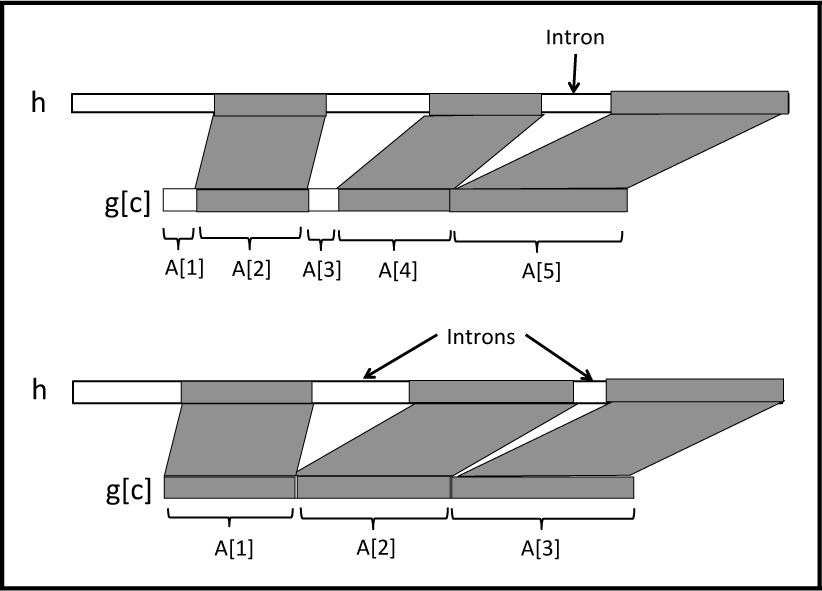
**Top.** Illustration of a spliced alignment between a CDS sequence *g*[*c*] and a gene sequence *h*, composed of 5 blocks, 3 conserved blocks (*A*[2], *A*[4] and *A*[5]) and 2 deleted blocks (*A*[1] and *A*[3]). It induces 1 putative intron between the successive conserved blocks *A*[4] and *A*[5]. The deleted blocks *A*[1] = (*k*_1_, *l*_1_, *a*_1_, *b*_1_) and *A*[3] = (*k*_3_, *l*_3_, *a*_3_, *b*_3_) are such that *a*_1_ = *b*_1_ = 0 and *a*_3_ = *b*_3_ = 0. **Bottom.** A spliced alignment composed of 3 conserved blocks that induce 2 putative introns between *A*[1] and *A*[2] and between *A*[2] and *A*[3].

Condition 1. of the spliced alignment definition implies that the set of conserved and deleted blocks of the spliced alignment covers the entire CDS sequence. Condition 2. implies that the blocks are ordered in the alignment following an increasing order of their location on the CDS sequence, and this order is also preserved on the gene sequence.

For instance, in Figure 9, the spliced alignment of *g*[*c*_1_] against *g* is *A* = {(1, 4, 9, 12), (5, 16, 18, 29), (17, 24, 49, 56)} (see Figure 10 for more general examples of spliced alignments with conserved and deleted blocks).

A spliced alignment *A* induces a set of putative gene intron segments. These intron segments are the gene segments that lie between two successive blocks of the spliced alignment that are conserved blocks. Formally, the set of introns induced by a spliced alignment *A* =*{*(*k*_1_, *l*_1_, *a*_1_, *b*_1_), …, (*k*_*j*_, *l*_*j*_, *a*_*j*_, *b*_*j*_)*}* is denoted by Intron(*A*) =*{*(*b*_*i*_, *a*_*i*+1_) *such that* (*k*_*i*_, *l*_*i*_, *a*_*i*_, *b*_*i*_) *and* (*k*_*i*+1_, *l*_*i*+1_, *a*_*i*+1_, *b*_*i*+1_) *are conserved blocks}*. Two successive conserved blocks of the spliced alignment *A* also induce a junction between two successive segments in the CDS sequence that are separated by an intron segment in the gene sequence. The set of putative exon junctions induced the spliced alignment *A* is denoted by Junction(*A*) = *{l*_*i*_ *such that* (*k*_*i*_, *l*_*i*_, *a*_*i*_, *b*_*i*_) *and* (*k*_*i*+1_, *l*_*i*+1_, *a*_*i*+1_, *b*_*i*+1_) *are conserved blocks}*. Note that if all blocks composing *A* are conserved, then the number of introns induced by *A* is |Intron(*A*)| = |Junction(*A*)| = |*A| -* 1.

### 3.2 A new constrained version of the spliced alignment problem

In this subsection, we first re-call two existing versions of the spliced alignment problem and we introduce a third more constrained version that takes additionally account of the splicing structure of input sequences. We discuss the motivations and limits of the different versions as we give their definitions.

Given a block (*k, l, a, b*) of a spliced alignment of a CDS sequence *g*[*c*] against a gene sequence *h*, let sim(*g*[*c*][*k, l*], *h*[*a, b*]) denote the score of an optimal global alignment between the CDS segment *g*[*c*][*k, l*] and the gene segment *h*[*a, b*] in a given scoring scheme. Note that if *a* = *b* = 0, then *h*[*a, b*] is an empty segment. The following is a reformulation of the less constrained version of the spliced alignment problem introduced in [8]. This formulation gives a unified framework to formally define and compare all the versions of the spliced alignment problem formulated thereafter.

#### Spliced Alignment Problem I (SAP_I)

**Input:** A CDS sequence *g*[*c*] from a gene sequence *g*; a gene sequence *h*.

**Output:** A spliced alignment *A* of *g*[*c*] against *h* that maximizes

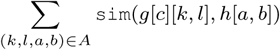

The SAP_I problem accounts only for the sequence similarity between the segments composing the blocks of the spliced alignment. In practice, in more than 99% of real cases of splicing, the spliced intron sequences start with a dinucleotide sequence “GT” and ends with a dinucleotide sequence “AG” corresponding to *canonical splice sites* [4, 20]. They exists also other non-canonical splice site pairs such as GC-AG, AT-AC that occur less frequently. Thus, in order to improve the accuracy of spliced alignments, a more constrained version of the problem allows to account for the extremities of intron segments induced by a spliced alignment. Given an intron (*b, a*) induced by two successive conserved blocks of a spliced alignment of a CDS sequence *g*[*c*] against a gene sequence *h*, let `splicesignals`(*h*[*b, a*]) denote a score of the putative intron segment *h*[*b, a*] accounting for the presence or absence of known splice signals at the extremities of *h*[*b, a*]. A putative intron segment with two canonical splice signals at its extremities has a higher score than a segment with only one which has a higher score than a segment without any canonical splice signal at its extremities. A more constrained version of the spliced alignment problem studied in [12, 25] for instance is the following.

#### Spliced Alignment Problem II (SAP_II)

**Input:** A CDS sequence *g*[*c*] from a gene sequence *g*; a gene sequence *h*.

**Output:** A spliced alignment *A* of *g*[*c*] against *h* that maximizes

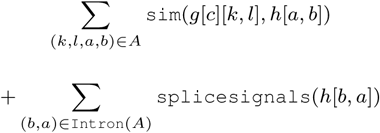

The SAP_I and SAP_II problems do not account for the splicing structure of the input sequences. In order to further improve the accuracy of spliced alignments, we define a more constrained version of the problem that takes into account the exon structure of the CDS sequence and the exon-intron structure of the gene sequence.

Given a putative intron (*b, a*) *∈* `Intron`(*A*) induced by two successive conserved blocks of a spliced alignment *A* of a CDS sequence *g*[*c*] against a gene sequence *h*, let `splicesites` *ε* (*h*)(*b, a*) denote a score of the putative intron segment *h*[*b, a*] accounting for the correspondance of its extremities with known splice sites in the gene sequence *h*. A putative intron segment whose extremities both correspond to known splice sites in the gene sequence receives a higher score than a segment with only one extremity corresponding to a known splice site which has a higher score than a segment without any correspondance to known splice sites at its extremities.

Similarly, for a putative CDS exon junction *l ∈* Junction(*A*) induced by two successive conserved blocks of the spliced alignment *A*, let exonjunction*∈* _(*g*[*c*])_(*l*) denote a score of the putative CDS `exon junction` *l* accounting for its correspondance with a real exon junction in the CDS sequence *g*[*c*]. If *l* corresponds to a real exon junction in *∈* (*g*[*c*]) it receives a higher score than if it does not correspond to a junction in *∈* (*g*[*c*]). The more constrained version of the problem introduced here is defined as follows.

#### Spliced alignment Problem III (SAP_III)

**Input:** A CDS sequence *g*[*c*] from a gene sequence *g*; the set of exons *ε* (*g*[*c*]) of *g*[*c*]; a gene sequence *h*; the set of exons *ε* (*h*) of *h*.

**Output:** A spliced alignment *A* of *g*[*c*] against *h* that maximizes

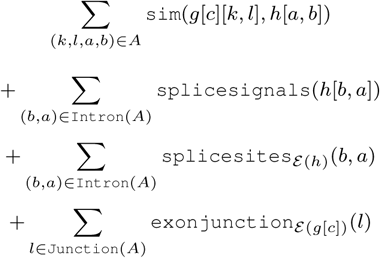

Examples of algorithms developed for the SAP_I problem which accounts only for the sequence similarity at the nucleotide or at the amino acid level are BLAT [14], DDS/GAP2 [10], SOAPsplice [9] and HSA [3]. Several algorithms have also been developed for the SAP_II problem, for instance Geneseqer [22], Sipdey [24], Splign [12], SIM4 [7], MGAlign [18], GMAP [25] and Spa [23]. These algorithms account for splice signals in addition to sequence similarity. A comparison of a subset of these tools has demonstrated the superiority of splice signal-based methods compared to only sequence similarity-based methods [25]. Moreover, it was shown in [12] that, among splice signal-based methods, the best performing current spliced alignment method for cDNA-to-gene alignment was Splign. Thus, taking account of more information about the input sequences improves the accuracy of spliced alignments. We then expect that accounting for information on the exon structure of CDS sequences and the exon-intron structure of gene sequences will further improve spliced alignments.

### 3.3 The SFA-align algorithm for the SAP_III problem

In this section, we describe a heuristic algorithm called SFA-align for the SAP_III problem. The algorithm follows a general approach followed by most heuristic methods developed for the SAP_I and SAP_II problems. The algorithm is decomposed into three steps that each accounts for the splicing structure of the input sequences. Note that in the case where the splicing structures of the input sequences are not provided, SFA-align includes a preliminary step in which the splicing structure of input sequences are inferred by computing a spliced alignment of each CDS against its own gene having a Percent Sequence Identity (PID) of 100. An overview of the main steps of SFA-align is depicted in Figure 1. It starts with a fast local alignment to compute highly conserved local segments used as anchors in the alignment. Next, the anchor alignment are extended in order to maximize the exon coverage. Finally, between the anchors, a global alignment of the remaining segments can be applied to complete the alignment.

**Step 1. Local alignments using tblastx.** This step is achieved using Translated Blast (tblastx) [13] in order to obtain a preliminary set of local alignments between the input CDS sequence and gene sequence. Tblastx is used in order to account for the translation of the sequences into amino acid sequences. This allows to detect conserved exon segments translated into amino acid sequences even in the presence of translational frameshifts or nucleotide silent mutations.

Given the set of hits obtained using tblastx with a given threshold E-value, the following procedure is applied to obtain the final set of local alignments. i) Each hit is assigned to the exon of the CDS that the hit covers the more, and the hit is minimally trimmed at its extremities to cover only this CDS exon. Thus, the boundaries of a trimmed hit never exceed the boundaries of the CDS exon that it is assigned to; ii) All hits within the same CDS exon are compared and only a subset of pairwise compatible hits with the lowest E-values is kept; iii) Finally, the hits within different exons are compared and only compatible hits with the lowest E-values are kept; iv) For each exon, the remaining compatible hits are gathered into a set of non-overlapping local alignments for the exon. A more detailed description of this procedure is given in Algorithm 1 in Appendix. Figure 11 provides an illustration of the result of the procedure on an example. In this example, Step i of the local alignment procedure associates hits 1 and 2 to exon *E*_1_, hits 3 and 4 to exon *E*_2_, hit 5 to exon *E*_3_, and hit 6 to exon *E*_4_. The extremities of the hit 5 are also trimmed so that the hit covers only exon *E*_3_. Step ii removes hit 4 from the list of hits associated to exon *E*_2_ because it is incompatible with hit 3 that has a lower E-value. Next, Step iii removes hit 6 from the list of hits associated to exon *E*_4_ because it is incompatible with hit 5 associated to exon *E*_3_ with a lower E-value. Finally, Step iv ends up with three local alignments covering exons *E*_1_, *E*_2_ and *E*_3_ illustrated by hits 1, 3, 5 in Figure 12.

**Fig. 11:**
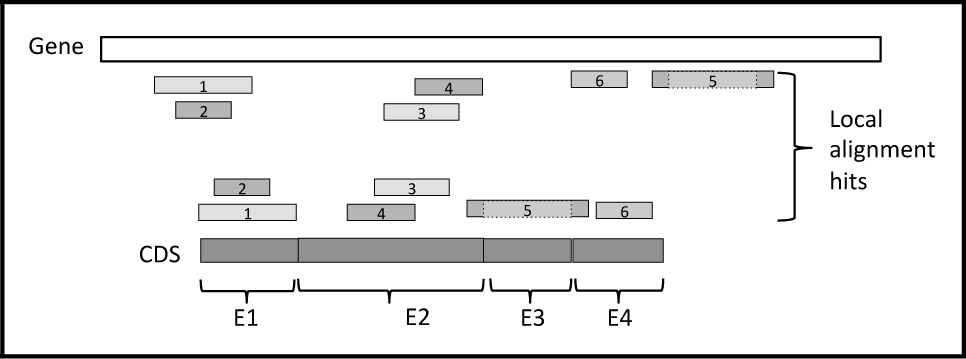
Example of a set of six hits obtained using tblastx between a CDS and a gene sequence, with hits numbered from 1 to 6 by increasing E-values. The local alignment algorithm ends up with the three local alignments 1, 3, 5 covering exons *E*_1_, *E*_2_ and *E*_3_.

**Fig. 12:**
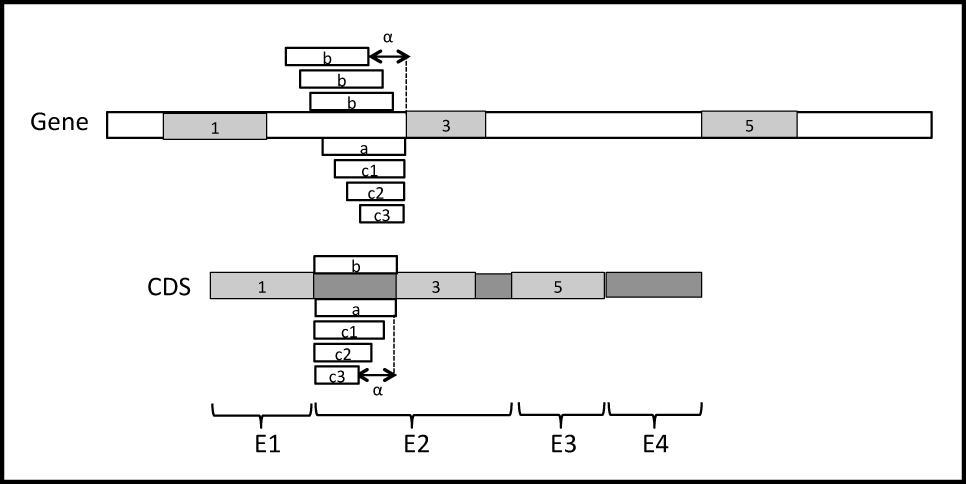
Example of a set of three local alignments 1, 3, 5 between a CDS and a gene sequence obtained at the end of the local alignment step. All possible configurations (a), (b) and (c) of left extension of the local alignments 3 are illustrated. The maximum number of gaps *α* for configurations (b) and (c) is also illustrated.

**Step 2. Gapped extension of anchors.** In this step, each local alignment obtained from Step 1 is extended in both directions in order to increase as much as possible the coverage of the CDS exon to which it is associated. The extension procedure allows an extended portion of an alignment to start with a succession of gaps whose number must be a multiple of 3 and shall not exceed a given number *α* of gaps. For example, a local alignment (“AAUCGGA”,“AAUCGGA”) that partially covers a CDS exon “AAUCGGAUGGGUG” could be extended on the right until the extremity of the exon following three possible configurations. It can be extended as (“AAUCGGAUGGGUG”,“AAUCGGAUGGGUG”) without any gap at the start of the extension, or (b) (“AAUCGGAUGGGUG”,“AAUCGGA---GUG”) starting with 3 gaps in the gene sequence, or (c) (“AAUCGGA---UGGGUG”, “AAUCGGACCCUGGGUG”) starting with 3 gaps in the CDS.

Given a local alignment of a CDS segment and a gene segment represented by a block (*k, l, a, b*) such that *k* and *l* are the start and end positions of the segment in the CDS, and *a* and *b* are the start and end positions of the segment in the gene sequence, the gapped extension procedure applied on (*k, l, a, b*) is as follows. Let (*k′*^*t′*^, *l*^*t*^) be the exon of the CDS to which the local alignment (*k, l, a, b*) is associated. Note that *k*^*t*^ ≤ *k* ≤ *l* ≤ *l*^*t*^. i) If *k*^*t*^ *< k*, then the alignment (*k, l, a, b*) can be extended on the left. The procedure tries all possible configurations of extension, (a) without any gap at the start of the extension, (b) starting with gaps in the gene sequence or (c) starting with gaps in the CDS, such that the extension does not overlap any other local alignment. For configurations and (c), all possible numbers of gaps 3 *\* i* with *i* ranging from 1 and *α*/3 are evaluated, the extension configuration having the highest identity score is returned. If this identity score is above a given identity threshold *β*, then the alignment is extended on the left, otherwise no extension is applied. i) If *l* < *l*^*t*^, then the alignment (*k, l, a, b*) can also be extended on the right. A procedure similar to the previous one is applied in order to try all possible configurations of extension on the right. A more detailed description of the procedure is given in Algorithm 2 in Appendix. Figure 12 also provides an illustration of the configurations of left extension that are explored by the procedure on an example.

**Step 3. Global spliced alignment algorithm.** For all CDS exons that were left completely unaligned by the previous steps, a dynamic programming algorithm for global spliced alignment is applied. Restricting the global alignment to remaining unaligned exons of the CDS allows to accelerate the global algorithm step by dividing the dynamic programming space. The following main recurrence formula is used.

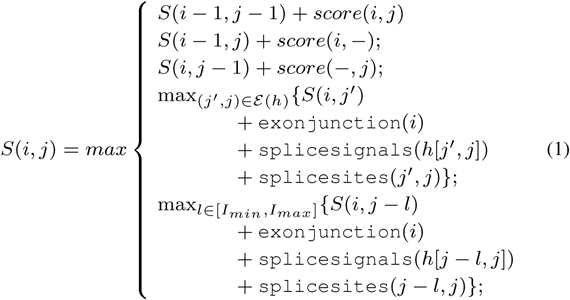

In Formula (1), when computing the global spliced alignment between a CDS segment *g*[*c*][*k, l*] and a gene segment *h*[*a, b*], *S*(*i, j*) is the maximum score of a global spliced alignment of the segment *g*[*c*][*k, k* + *i -* 1] and the segment *h*[*a, a* + *i -* 1]. For instance, if *g*[*c*][*k, l*] has length *m* and *h*[*a, b*] has length *n*, then *S*(*m, n*) is the maximum score of a global spliced alignment of *g*[*c*][*k, l*] and the segment *h*[*a, b*].

In the formula, the three first cases contribute to the computation of sequence alignment scores within blocks of the spliced alignment, given *score*(*i, j*), *score*(*i, -*), *score*(*-, j*) that denote the scores of substitution, insertion and deletion of nucleotides. The two last cases contribute to evaluating the structure alignment score according to the correspondence of induced introns with known splicing signals and the splicing structure of input sequences. Formula (1) is an extension of the main recurrence formula used in the global alignment step of Splign [12]. The extension consists in accounting for exonjunction(*i*) and splicesites(*j′, j*) that are the scores for CDS exon junctions and genomic introns introduced for the definition of the SAP_III problem. The last case of the formula also contributes to the extension and allows to further account for the splicing structure of the unspliced genomic sequence.

Finally, any new alignment block computed in this step is kept if and only if its identity score is above a given identity threshold *β*, otherwise it is not added to spliced alignment.

### 3.4 The SFA-ortholog algorithm for the identification of CDS orthology groups

In this section, we give a definition of CDS orthology groups computed by the SFA-ortholog algorithm based on pairwise spliced alignments. We first start with a definition of orthologous CDS based on spliced alignment.

In [26], an extension of the concept of gene orthology to spliced transcript orthology was introduced. They defined orthologous transcripts as two structurally similar transcripts from two orthologous genes. The orthology relationship between two transcripts relies on the structural similarity between the transcripts. This structural similarity is evaluated using the CDS associated to the transcripts and their spliced alignments against genes.

**CDS orthology:** Let *c*_1_ and *c*_2_ be two CDS from two homologous genes *g* and *h* respectively. Let *A*_1_ be a spliced alignment of the CDS sequence *g*[*c*_1_] against the gene sequence *h*. The CDS *c*_1_ and *c*_2_ are orthologs if:

(1) *| c*_1_ | = *| c*_2_ |;

(2) Intron(*A*_1_) = Intron(*c*_2_);

(3) for any *i*, 1 *≤ i ≤ | c*_1_ |, [length(*c*_1_[*i*])-length(*c*_2_[*i*])] % 3 = 0.

In other terms, *c*_1_ are *c*_2_ as orthologs if (1) they have the same number of exons *| c*_1_ | = *| c*_2_ |; (2) the spliced alignment of *c*_1_ against *h* induces the same introns for *c*_1_ and *c*_2_, Intron(*A*_1_) = Intron(*c*_2_); (3) the lengths of each pair of corresponding exons in *c*_1_ and *c*_2_ are congruent modulo 3. Conditions (1) and (2) ensure that the two CDS have the same splicing structure. Condition (3) ensures that the two CDS are translated in the same codon phase in each pair of corresponding exons in order to generate similar protein sequences.

Note that this definition only requires that one of the spliced alignments of *g*[*c*_1_] against *h* or *h*[*c*_2_] against *g* supports the orthology relation. An alternative more stringent definition of CDS orthology consists in requiring the reciprocity, i.e. that both spliced alignment support the orthology relation.

**CDS orthology groups:** Given a set of CDS *C* from a set of homologous genes *G*, the transitivity of the CDS orthology relation is assumed and used to identify distant orthologs in *C* that cannot be directlty identified by means of the CDS structural similarity. Such orthologs could be missed because of partial spliced alignments due to low sequence similarity.

The CDS orthology relation on *C* is then extended into an equivalence relation such that for any three CDS *c*_1_, *c*_2_, *c*_3_ in *C*, if *c*_1_ and *c*_2_ are orthologs and *c*_2_ and *c*_3_ are orthologs, then *c*_1_ and *c*_3_ are also orthologs. The CDS orthology groups are defined as the equivalence classes of the resulting equivalence relation.

## 4. Conclusion

The article introduces a new version of the spliced alignment problem accounting for the splicing structure of input sequences. It constitutes a new approach to compute accurate CDS-to-gene spliced alignments, by detecting conservations in the splicing structure of input sequences.

We present a heuristic algorithm for the problem, and we show that it is useful to improve the accuracy of spliced alignments and to identify CDS orthology groups within a set of CDS from homologous genes. The application of the algorithm to real and simulated datasets shows that the new method outperforms existing spliced alignment methods in terms of accuracy, with comparable execution times for CDS-to-gene spliced alignment, and its performance is robust to changes in the level of input sequence similarity.

## Appendix

### Algorithm 1 Local alignment

~~~
**INPUT:** *list*_*of* _*hits*: list of tblastx hits
**OUTPUT:** *list*_*of* _*anchors*: list of local alignments
**\*\*\*Step i)***
for** *H ∈ list*_*of* _*hits* **do**
  *E ←* find the exon of the CDS that the hit *H* covers the more
  *H ←* trim the hit *H* so that *H* covers only the exon *E hits*_*of* _*exon*[*E*] *←* add the hit *H* to *hits*_*of* _*exon*[*E*]
  **end for
***Step ii)***
for** *E ∈ CDS*_*exons* **do**
  *kept*_*hits*_*of* _*exon*[*E*] *←*[]
  *hits*_*of* _*exon*[*E*] *←* sort *hits*_*of* _*exon*[*E*] by increasing E-value
  **for** *H ∈ hits*_*of* _*exon*[*E*] **do
     if** H is compatible with all hits in *kept*_*hits*_*of* _*exons*[*E*] **then**
       *kept*_*hits*_*of* _*exon*[*E*] *←* add *H* to *kept*_*hits*_*of* _*exon*[*E*]
     **end if
  end for
end for
***Step iii)***
for** (*E*_1_, *E*_2_) *∈ CDS*_*exons*^2^ **do
  for** (*H*_1_, *H*_2_) *∈ kept*_*hits*_*of* _*exon*[*E*_1_] ×
    *kept*_*hits*_*of* _*exon*[*E*_2_] **do
      if** *H*1 and *H*2 are not compatible **then**
         keep the hit with the lower E-value and discard the other
    **end if
  end for
end for
***Step iv)***
for** *E ∈ CDS*_*exons* **do**
  *S ←* merge *hits*_*of* _*exon*[*E*] into a set *S* of non-overlapping local alignments (*k, l, a, b*) of CDS segments (*k, l*) and gene segments
  (*a, b*)
  *list*_*of* _*anchors ←* add set *S* of local alignments to
  *list*_*of* _*anchors*
**end for**
~~~

### Algorithm 2 Gapped extension

~~~
**INPUT:** *list*_*of* _*anchors*: list of local alignments
**OUTPUT:** *list*_*of* _*ext*_*anchors*: list of extended local alignments
**for** (*k, l, a, b*) *∈ list*_*of*_*anchors* **do**
  (*k′, l′*) *←* CDS exon covered by (*k, l, a, b*)
  **\*\*\*Step i) Extension on the left***
  if** *k′ < k* **then
      ***Extension configuration (a)\*\*\***
      *max*_*identity*_*extension ←* (*k′, k, a -* (*k - k′*), *a*)
      *max*_*identity ← PID*(*max*_*identity*_*extension*)
      **\*\*\*Extension configuration (b)***
      for** 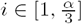 **do**
        *extension ←* (*k′, k, a* - (*k - k′*) - 3*i, a* - 3*i*)
        *identity*_*extension ← PID*(*extension*)
        **if** *identity*_*extension > max*_*identity* **then**
          *max*_*identity*_*extension ← extension
          max*_*identity ← identity*_*extension*
        **end if
  end for
  ***Extension configuration (c)***
  for** 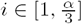 **do**
    *extension ←* (*k′, k -* 3*i, a -* (*k -* 3*i - k′*), *a*)
    *identity*_*extension ← PID*(*extension*)
    **if** *identity*_*extension > max*_*identity* **then**
      *max*_*identity*_*extension ← extension
      max*_*identity ← identity*_*extension*
    **end if
  end for
  if** *max*_*identity = β* **then**
    *left*_*extension ← extension*
  **end if
end if
  ***Step ii) Extension on the right***
  if** *l < l′* **then**
    *right*_*extension ←* procedure symmetric to the one used in Step
    i) for extension on the left
  **end if
end for**
(*k, l, a, b*) *←* merge *left*_*extension*, (*k, l, a, b*) and *right*_*extension* into a single extended alignment
*list*_*of* _*ext*_*anchors ←* add (*k, l, a, b*) in *list*_*of* _*ext*_*anchors*
~~~

## References

[1] Samuel Blanquart, Jean-Stéphane Varré, Paul Guertin, Amandine Perrin, Anne Bergeron, and Krister M Swenson. Assisted transcriptome reconstruction and splicing orthology. BMC Genomics, 17(10):157, 2016.

[2] Volker Brendel, Liqun Xing, and Wei Zhu. Gene structure prediction from consensus spliced alignment of multiple ests matching the same genomic locus. Bioinformatics, 20(7):1157–1169, 2004.

[3] Jingde Bu, Xuebin Chi, and Zhong Jin. Hsa: a heuristic splice alignment tool. BMC systems biology, 7(2):S10, 2013.

[4] M Burset, IA Seledtsov, and VV Solovyev. Analysis of canonical and non-canonical splice sites in mammalian genomes. Nucleic Acids Research, 28(21):4364–4375, 2000.

[5] Yann Christinat and Bernard ME Moret. Inferring transcript phylogenies. BMC Bioinformatics, 13(9):S1, 2012.

[6] Fiona Cunningham, M Ridwan Amode, Daniel Barrell, et al. Ensembl 2015. Nucleic Acids Research, 43(D1):D662–D669, 2015.

[7] Liliana Florea, George Hartzell, Zheng Zhang, Gerald M Rubin, and Webb Miller. A computer program for aligning a cdna sequence with a genomic dna sequence. Genome research, 8(9):967–974, 1998.

[8] Mikhail S. Gelfand, Andrey A. Mironov, and Pavel A. Pevzner. Spliced alignment: A new approach to gene recognition, pp. 141–158. Springer Berlin Heidelberg, Berlin, Heidelberg, 1996.

[9] Songbo Huang, Jinbo Zhang, Ruiqiang Li, Wenqian Zhang, Zengquan He, Tak-Wah Lam, Zhiyu Peng, and Siu-Ming Yiu. Soapsplice: genome-wide ab initio detection of splice junctions from rna-seq data. Frontiers in genetics, 2:46, 2011.

[10] Xiaoqiu Huang, Mark D Adams, Hao Zhou, and Anthony R Kerlavage. A tool for analyzing and annotating genomic sequences. Genomics, 46(1):37–45, 1997.

[11] Zhengyan Kan, Eric C Rouchka, Warren R Gish, et al. Gene structure prediction and alternative splicing analysis using genomically aligned ests. Genome Research, 11(5):889–900, 2001.

[12] Yuri Kapustin, Alexander Souvorov, Tatiana Tatusova, and David Lipman. Splign: algorithms for computing spliced alignments with identification of paralogs. Biology direct, 3(1):20, 2008.

[13] Harpreet Kaur, Amandeep Singh, and Pardeep Singh. Comparison of variants of blast. In Proceedings of the International MultiConference of Engineers and Computer Scientists, volume 1, 2008.

[14] W James Kent. BlatâŁ“the blast-like alignment tool. Genome research, 12(4):656–664, 2002.

[15] Rodrigo Mitsuo Kishi, Ronaldo Fiorilo dos Santos, and Said Sadique Adi. Gene prediction by multiple spliced alignment. In Brazilian Symposium on Bioinformatics, pp. 26–33. Springer, 2011.

[16] Carolin Kosiol, Ian Holmes, and Nick Goldman. An empirical codon model for protein sequence evolution. Molecular biology and evolution, 24(7):1464–1479, 2007.

[17] Esaie Kuitche, Manuel Lafond, and Aïda Ouangraoua. Reconstructing protein and gene phylogenies by extending the framework of reconciliation. Proceedings of International Conference on Bioinformatics and Computational Biology (BICOB’17), (ISBN:9781510836679):79–86, 2017.

[18] Shoba Ranganathan, Bernett TK Lee, and Tin Wee Tan. Mgalign, a reduced search space approach to the alignment of mrna sequences to genomic sequences. Genome Informatics, 14:474–475, 2003.

[19] Jose Manuel Rodriguez, Paolo Maietta, Iakes Ezkurdia, Alessandro Pietrelli, Jan-Jaap Wesselink, Gonzalo Lopez, Alfonso Valencia, and Michael L Tress. Appris: annotation of principal and alternative splice isoforms. Nucleic acids research, 41(D1):D110–D117, 2012.

[20] Michael E Sparks and Volker Brendel. Incorporation of splice site probability models for non-canonical introns improves gene structure prediction in plants. Bioinformatics, 21(Suppl_3):iii20–iii30, 2005.

[21] Mario Stanke, Ana Tzvetkova, and Burkhard Morgenstern. Augustus at egasp: using est, protein and genomic alignments for improved gene prediction in the human genome. Genome biology, 7(1):S11, 2006.

[22] Jonathan Usuka, Wei Zhu, and Volker Brendel. Optimal spliced alignment of homologous cdna to a genomic dna template. Bioinformatics, 16(3):203–211, 2000.

[23] Erik Van Nimwegen, Nicodeme Paul, Robert Sheridan, and Mihaela Zavolan. Spa: a probabilistic algorithm for spliced alignment. PLoS genetics, 2(4):e24, 2006.

[24] Sarah J Wheelan, Deanna M Church, and James M Ostell. Spidey: a tool for mrna-to-genomic alignments. Genome research, 11(11):1952–1957, 2001.

[25] Thomas D Wu and Colin K Watanabe. Gmap: a genomic mapping and alignment program for mrna and est sequences. Bioinformatics, 21(9):1859–1875, 2005.

[26] Federico Zambelli, Giulio Pavesi, Carmela Gissi, David S Horner, and Graziano Pesole. Assessment of orthologous splicing isoforms in human and mouse orthologous genes. BMC Genomics, 11(1):1, 2010.

